# Is virulence phenotype evolution driven exclusively by *Lr* gene deployment in French *Puccinia triticina* populations?

**DOI:** 10.1101/2022.03.16.484559

**Authors:** Cécilia Fontyn, Anne-Catherine Zippert, Ghislain Delestre, Thierry C Marcel, Frédéric Suffert, Henriette Goyeau

**Affiliations:** Université Paris-Saclay, INRAE, UR BIOGER, 78850 Thiverval-Grignon, France

**Keywords:** Aggressiveness, leaf rust, *Lr* gene, pathotype distribution, *Puccinia triticina*, varietal landscape, virulence, wheat pathogen

## Abstract

*Puccinia triticina* is a highly damaging wheat pathogen. The efficacy of leaf rust control by genetic resistance is mitigated by the adaptive capacity of the pathogen, expressed as changes in its virulence combinations (pathotypes). An extensive *P. triticina* population survey has been carried out in France over the last 30 years, describing the evolution dynamics of this pathogen in response to cultivar deployment. We analyzed the dataset for the 2006-2016 period to determine the relationship between the *Lr* genes in the cultivars and virulence in the pathotypes. Rust populations were dominated by a small number of pathotypes, with variations in most of the virulence frequencies related to the corresponding *Lr* gene frequencies in the cultivated landscape. Furthermore, the emergence and spread of a new virulence matched the introduction and use of the corresponding *Lr* gene (*Lr28*), confirming that the deployment of qualitative resistance genes is an essential driver of evolution in *P. triticina* populations. However, PCA revealed that certain pathotype-cultivar associations cannot be explained solely by the distribution of *Lr* genes in the landscape. This conclusion is supported by the predominance of a few pathotypes on some cultivars, with the persistence of several other compatible pathotypes at low frequencies. Specific interactions are not, therefore, sufficient to explain the distribution of virulence in rust populations. Our findings suggest that aggressiveness is a driver of changes in pathotype frequencies. Accordingly, the hypothesis of “dual selection”, based on both qualitative and quantitative interactions between *P. triticina* populations and bread wheat cultivars, is favored.

## INTRODUCTION

Leaf rust caused by *Puccinia triticina* is one of the most damaging wheat diseases, with high yield losses at worldwide (Huerta-Espino et al. 2011; Savary et al. 2019). This pathogen is a heteroecious biotrophic fungus that can infect bread wheat (*Triticum aestivum*) and durum wheat (*Triticum durum*) as primary hosts. *P. triticina* requires an alternative host, *Thalictrum speciosissimum*, for its sexual reproduction and, thus, for completion of its life cycle (Bolton et al. 2008). The alternative host is not naturally present in most places worldwide (including France), and *P. triticina* is, therefore, generally found exclusively in its uredinial and telial stages on wheat (Kolmer 2013). The proportion of repeated genotypes is high and there is an excess of heterozygotes in European and French leaf rust populations, confirming the principal role of clonal reproduction (Goyeau et al. 2007; Kolmer et al. 2013).

Genetic resistance is the cheapest and most effective means of limiting leaf rust epidemics. Eighty *Lr* genes, most displaying gene-for-gene (qualitative) interactions, have been identified in wheat cultivars. The gene-for-gene relationship depends on the presence of a virulence (*Avr*) gene in the pathogen (Flor 1971; Van Der Biezen & Jones 1998). A virulence phenotype, also called “pathotype” or “race”, is defined by a virulence profile: two pathogenic strains are considered to belong to the same pathotype if they have the same combination of virulence genes. Some *Lr* genes express a partial type of resistance (quantitative or partial) at the adult plant stage (McIntosh et al. 2017; Prasad et al. 2020). A large proportion of these adult plant resistance (APR) genes, which may have a minor to intermediate effect, appear to be “race non-specific”, i.e. their effect is independent of the virulence phenotype of the pathogen (Lagudah 2011). Such resistance reduces disease symptoms, as shown, for example for *Lr34*, which increases the latency period and decreases the production of uredinia (Drijepondt 1989), or for *Lr46, Lr67* and *Lr68* (Singh et al. 1998; Herrera-Foessel et al. 2011; 2012), which increase the latency period and result in the production of fewer, smaller pustules (Rosewarne et al. 2006; Huerta-Espino et al. 2020).

Host genetic resistance can be broken down or eroded as a result of the evolution of pathogen populations. Indeed, despite their clonality, *P. triticina* populations are characterized by high diversity, achieved through a combination of asexual reproduction, mutation, and migration between wheat-growing areas via the long-distance dispersal of urediniospores. Somatic exchanges in *P. triticina* can also be a significant source of diversity and allelic rearrangements leading to new combinations of virulence genes (Figueroa et al. 2020). The evolution of *P. triticina* populations is heavily influenced by gene-for-gene interactions, expressed as “boom-and-bust” cycles of resistance (McDonald & Linde 2002). Typically, a wheat cultivar with a new single resistance gene is introduced widely into the landscape (“boom”), and the selection pressure imposed by this resistant cultivar then leads to adaptation of the pathogen population, by mutation, from avirulence to virulence. The new resistance gene loses efficacy as the new virulent population of the pathogen increases (“bust”) (McIntosh & Brown 1997; McDonald & Linde 2002). In this way, the pathogen has developed virulence toward most of the existing *Lr* genes, but the frequencies and combinations of virulence genes in pathogen populations vary over space and time (Park et al. 2002; Bolton et al. 2008; Huerta-Espino et al. 2011; Kolmer 2013, 2019).

The adaptation dynamics of *P. triticina* populations within the varietal landscape result in a rapid, continuous turnover of the predominant virulence phenotypes. Surveys of both host and pathogen populations in the main wheat-growing areas are, therefore, essential to increase the efficacy and durability of genetic resistances. Leaf rust populations have been monitored for a number of years, through field surveys performed around the world (Young, Jr. 1977; Walid et al. 2015; Kosman et al. 2019; Prasad et al. 2017; Liu & Chen 2012; Gultyaeva et al. 2012). These surveys aim to evaluate changes in the prevalence and landscape distribution of virulence phenotypes, and to detect any new pathotypes (new virulences and combinations of virulences) that might pose a threat to wheat cultivars with effective leaf rust resistance genes. The datasets generated by these surveys are valuable resources that can help breeders and extension services to propose cultivar deployment strategies both to control leaf rust and to improve the durability of the resistance genes used. ‘Thatcher’ near-isogenic lines (NILs) were developed to standardize the comparison of virulences between *P. triticina* populations of different geographic origins (McIntosh et al. 1995) and have been used in diverse surveys around the world since 1993.

Most surveys have described only virulence combinations in *P. triticina* populations. Others have linked the virulences in the pathogen population to the *Lr* genes in the cultivars deployed, thereby highlighting the influence of cultivar landscape on the evolution of *P. triticina* populations at large spatiotemporal scales. Such surveys were conducted at the European scale for the 1996-1999 period (Mesterházy et al. 2000), and, more recently, from 2018 to 2021, for the European ‘RustWatch’ research program (http://rustwatch.au.dk/). Virulence frequencies mostly depend on the leaf rust resistance genes present in the most common varieties in the landscape, with similar results also obtained in the US (Kolmer 2019; Mesterházy et al. 2000). In France, the *Lr* gene content of newly released varieties is postulated each year, making it possible to provide a comprehensive and continuous snapshot of the frequency of these genes at landscape level. The *Lr* gene composition of two-thirds of the cultivars present in the French landscape under wheat from 1983 to 2007 has been analyzed, with the *Lr13* gene predominating, being found in 67% of cultivars (Goyeau & Lannou 2011). French leaf rust populations have been monitored for the last 30 years. The populations of strains collected over the 1999-2002 period had highly diverse *P. triticina* virulence phenotypes, with 104 different pathotypes identified (Goyeau et al. 2006). However, the survey revealed domination by a single pathotype (073 100 0) coinciding with a period during which the cultivar landscape was dominated by the cultivar ‘Soissons’ (up to 40% in 1993). Pathotype 073 100 0 was found to be more aggressive on this cultivar than other virulent pathotypes (Pariaud et al. 2009b). This adaptation to ‘Soissons’, probably due to the absence of an effective *Lr* gene, greater aggressiveness, and the high frequency of this cultivar in the landscape, led to the domination of pathotype 073 100 0 from 1999 to 2002. The French wheat landscape tended to diversify thereafter, and the frequency of pathotype 073 100 0 decreased following the decline of ‘Soissons’ in the landscape (Papaïx et al. 2011). The impact of host quantitative resistance on the evolution of the aggressiveness spectrum of pathogen populations is much less well documented than the impact of *Lr* genes on the prevalence of virulence phenotypes. However, it has been shown that fungal pathogens can evolve and adapt to quantitative resistance through selection for greater aggressiveness (Delmas et al. 2016; Frézal et al. 2018).

The objective of this study was to determine the impact of the overall composition of the varietal landscape on the evolution of *P. triticina* populations over a decade. To this end, we analyzed the dataset generated by the annual surveys performed over the 2006-2016 period to determine the prevalence of *P. triticina* virulence phenotypes (pathotypes) across France. The *Lr* genes in the most widely grown cultivars were postulated and we determined whether their distribution in the varietal landscape could account for the prevalence of certain pathotypes.

## MATERIALS AND METHODS

### Pathogen population sampling

*P. triticina* isolates were sampled annually during the 2006-2016 period from a network of nurseries not sprayed with fungicide at about 50 different sites in wheat-growing areas of France. Samples were collected by breeders (the French Wheat Breeders group ‘Recherches Génétiques Céréales’, CETAC) and extension services (ARVALIS-Institut du Végétal). The sampling effort focused on the most widely grown bread wheat cultivars, hereafter referred to as ‘major’ cultivars (i.e. 35 cultivars, each grown on at least 2% of the French wheat-growing area; Table 1), planted in small plots (10 to 20 m^2^) in all nurseries. The cumulative area under wheat planted with these cultivars accounted for between 42.9% and 56.0% of the total bread wheat landscape, depending on the year. For each plot, a few infected leaves of each major cultivar were collected in May or June of each year. A single pustule (uredinium) was selected and urediniospores were collected to obtain one isolate per cultivar and site. Leaf rust samples were also collected from ‘minor’ bread wheat cultivars (i.e. cultivars grown on less than 2% of the French wheat-growing area) growing in the same nurseries. These minor cultivars were not necessarily present at all sites. The sample obtained from minor cultivars was therefore generally smaller than the sample from major cultivars. Between 75% (2016) and 95% (2010) of the total number of isolates were collected from major cultivars (Table 2).

**Table 1.**
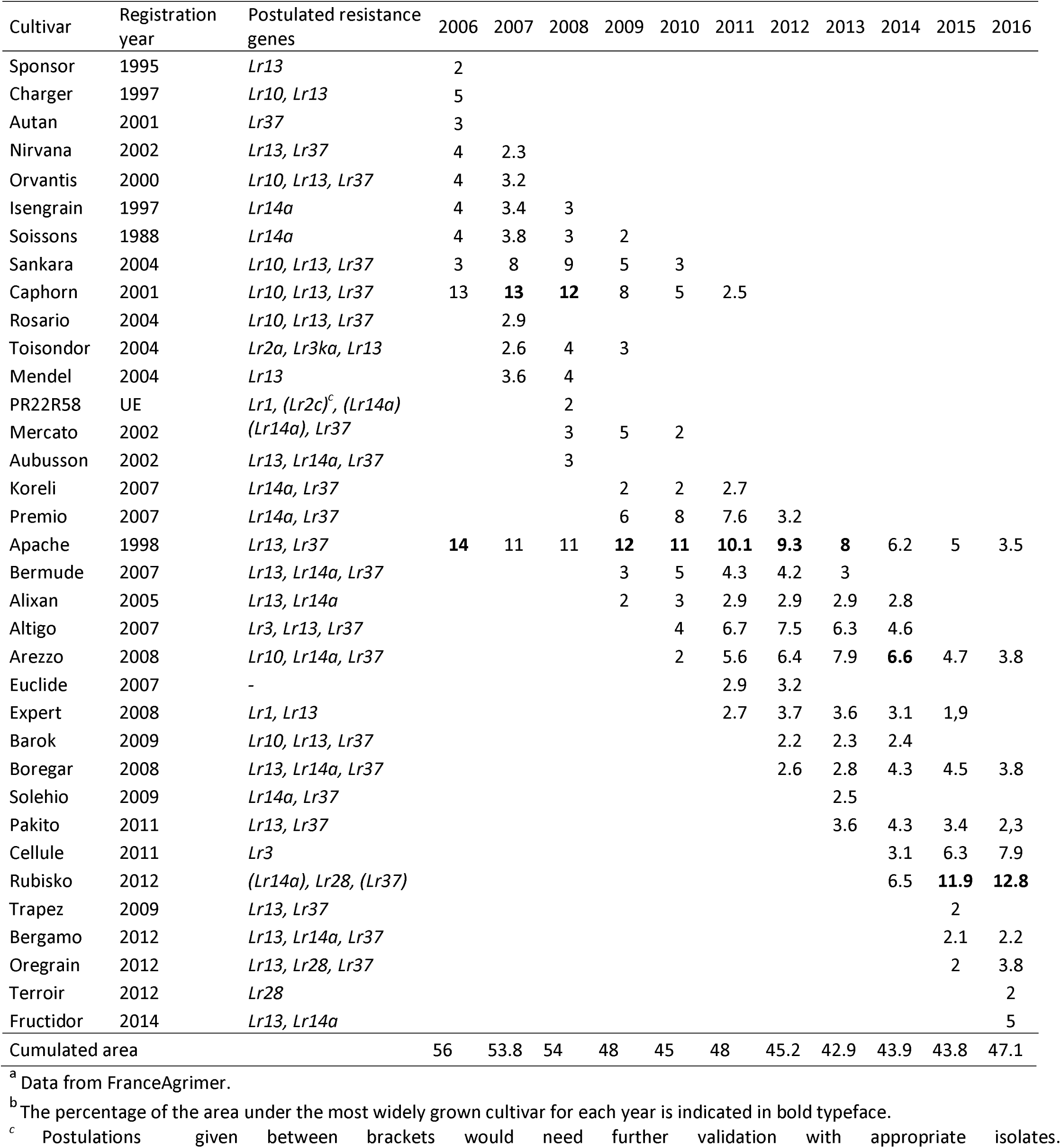
Bread wheat area (as a percentage) under various cultivars in France during the 2006-2016 period^a^. Only bread wheat cultivars accounting for at least 2% of the wheat-growing area for at least one year are indicated^b^, with their year of registration and postulated leaf rust resistance genes

**Table 2.** Number of isolates collected from the 35 major French wheat cultivars during the 2006-2016 period. Only bread wheat cultivars accounting for at least 2% of the wheat-growing area for at least one year are indicated, with their year of registration and postulated leaf rust resistance genes

| Cultivar | Registration year | Postulated resistance genes | 2006 | 2007 | 2008 | 2009 | 2010 | 2011 | 2012 | 2013 | 2014 | 2015 | 2016 |
| --- | --- | --- | --- | --- | --- | --- | --- | --- | --- | --- | --- | --- | --- |
| Sponsor | 1995 | *Lr13* |  |  |  |  |  |  |  |  |  |  |  |
| **Charger** | 1997 | *Lr10, Lr13* | 26 | 24 | 8 | 1 | 1 |  |  |  |  |  |  |
| Autan | 2001 | *Lr37* |  | 2 |  |  |  |  |  |  |  |  |  |
| Nirvana | 2002 | *Lr13, Lr37* |  | 1 | 1 |  |  |  |  |  |  |  |  |
| **Orvantis** | 2000 | *Lr10, Lr13, Lr37* | 39 | 48 | 16 | 1 | 2 | 1 |  |  |  |  |  |
| **Isengrain** | 1997 | *Lr14a* | 13 | 22 | 10 | 3 | 4 | 4 | 2 |  |  |  |  |
| **Soissons** | 1988 | *Lr14a* | 37 | 44 | 31 | 23 | 21 | 29 | 8 | 4 | 5 | 4 |  |
| **Sankara** | 2004 | *Lr10, Lr13, Lr37* | 10 | 21 | 18 | 18 | 6 | 6 | 1 |  |  |  |  |
| **Caphorn** | 2001 | *Lr10, Lr13, Lr37* | 45 | 64 | 41 | 33 | 27 | 18 | 1 |  |  |  |  |
| Rosario | 2004 | *Lr10, Lr13, Lr37* | 1 |  |  |  |  |  |  |  |  |  |  |
| Toison dor | 2004 | *Lr2a, Lr3ka, Lr13* |  |  | 2 |  |  | 2 |  |  |  |  |  |
| **Mendel** | 2004 | *Lr13* | 4 | 13 | 18 |  |  |  |  |  |  |  |  |
| PR22R58 | UE | *Lr1, (Lr2c)<sup>a</sup>, (Lr14a)* |  | 8 | 2 | 1 | 1 |  |  |  | 1 |  |  |
| Mercato | 2002 | *(Lr14a), Lr37* |  | 2 |  |  |  |  |  |  |  |  |  |
| **Aubusson** | 2002 | *Lr13, Lr14a, Lr37* | 11 | 28 | 18 | 23 | 29 | 33 | 21 | 1 |  |  |  |
| Koreli | 2007 | *Lr14a, Lr37* | 1 |  | 1 | 1 |  | 1 | 1 |  |  |  |  |
| **Premio** | 2007 | *Lr14a, Lr37* |  |  | 1 | 29 | 34 | 55 | 45 |  |  |  |  |
| Apache | 1998 | *Lr13, Lr37* | 47 | 71 | 43 | 44 | 38 | 62 | 50 | 39 | 46 | 45 | 21 |
| **Bermude** | 2007 | *Lr13, Lr14a, Lr37* |  |  |  |  | 16 | 38 | 35 | 24 | 31 | 30 |  |
| Alixan | 2005 | *Lr13, Lr14a* | 1 |  |  | 1 |  | 1 | 1 |  |  |  |  |
| **Altigo** | 2007 | *Lr3, Lr13, Lr37* |  | 1 |  |  | 2 | 3 | 35 | 26 | 11 |  |  |
| **Arezzo** | 2008 | *Lr10, Lr14a, Lr37* |  |  |  |  | 25 | 47 | 49 | 37 | 50 | 60 | 28 |
| Euclide | 2007 | - |  |  |  |  |  | 1 |  |  |  |  |  |
| **Expert** | 2008 | *Lr1, Lr13* |  |  |  |  |  | 2 | 21 | 16 | 16 | 16 | 5 |
| Barok | 2009 | *Lr10, Lr13, Lr37* |  |  |  |  | 1 |  |  |  |  |  |  |
| Boregar | 2008 | *Lr13, Lr14a, Lr37* |  |  |  |  |  |  |  |  |  |  |  |
| **Solehio** | 2009 | *Lr14a, Lr37* |  |  |  |  |  |  | 37 | 31 | 36 | 41 | 26 |
| **Pakito** | 2011 | *Lr13, Lr37* |  |  |  |  |  |  | 34 | 33 | 38 | 28 |  |
| **Cellule** | 2011 | *Lr3* |  |  |  |  |  |  | 2 | 5 | 45 | 38 | 35 |
| **Rubisko** | 2012 | *(Lr14a), Lr28, (Lr37)* |  |  |  |  |  |  | 1 | 7 | 14 | 34 | 18 |
| Trapez | 2009 | *Lr13, Lr37* |  |  |  |  |  |  |  |  |  | 1 |  |
| Bergamo | 2012 | *Lr13, Lr14a, Lr37* |  |  |  |  |  |  |  |  |  |  |  |
| **Oregrain** | 2012 | *Lr28* |  |  |  |  |  |  | 4 | 9 | 17 | 39 | 26 |
| Terroir | 2012 | *Lr28* |  |  |  |  |  |  |  |  | 13 | 17 | 8 |
| **Fructidor** | 2014 | *Lr13, Lr14a* |  |  |  |  |  |  |  |  |  | 1 | 16 |
| Others minor cultivars |  |  | 71 | 69 | 28 | 14 | 10 | 28 | 63 | 63 | 54 | 62 | 62 |
| Total |  |  | 306 | 418 | 238 | 192 | 217 | 331 | 411 | 295 | 377 | 416 | 245 |
| % of isolates collected on major wheat cultivars |  |  | 77 | 83 | 88 | 93 | 95 | 92 | 85 | 79 | 86 | 85 | 75 |

<sup>a</sup> Postulations given between brackets would need further validation with appropriate isolates.
Cultivars accounting for more than 2% of the pathogen samples in a given year are shown in bold typeface.

### Pathotype determination

The pathotype of each *P. triticina* isolate (i.e. its virulence combination) was determined *in planta*, by inoculating a differential set of wheat lines as described by Goyeau et al. (2011). Before inoculation, all healthy plant material was grown in cabinets with filtered air, in a glasshouse at temperatures between 15 and 20°C and with a 14-h photoperiod (daylight supplemented with light from 400 W sodium lamps). Infected leaves collected in the field were wiped gently on seven-day-old seedlings of the wheat cv. ‘Michigan Amber’ treated with 15 ml of maleic hydrazide solution (0.25 g of maleic hydrazide per liter of H_2_O) to prevent the emergence of secondary leaves and to increase the size of uredinia. Inoculated seedlings were placed for 24 h in a dew chamber at 15°C, then in the glasshouse. One week after inoculation, the seedlings were trimmed so that only one plant with one uredinium remained in each pot. Cellophane bags were then placed over the pots to prevent contamination between isolates. Spores from a single uredinium were then multiplied to produce batches of spores for storage and/or inoculation of a differential set of plant lines. To this end, the spores were collected in a gelatin capsule (size 00) with a cyclone collector 10 days after inoculation. The collected spores were then diluted by adding 0.5 ml of light mineral oil to each capsule, and the resulting spore suspension was sprayed onto seven-day-old ‘Michigan Amber’ seedlings. The differential sets were sown in pressed-peat pots (3 × 3 cm2) containing a commercial compost (peat substrate; Gebr. Brill Substrate, Georgsdorf, Germany), with four seedlings per pot and two pots per line (eight seedlings for each differential line). The differential set consisted of 20 lines (18 ‘Thatcher’ NILs carrying *Lr1, Lr2a, Lr2b, Lr2c, Lr3a, Lr3bg, Lr3ka, Lr10, Lr13, Lr14a, Lr15, Lr16, Lr17, Lr20, Lr23, Lr24, Lr26*, and *Lr37*, ‘CS2A/2M’ carrying *Lr28*, the Australian wheat cv. ‘Harrier’ carrying *Lr17b*; Singh et al. 2001) and the susceptible wheat cultivar ‘Morocco’ as a control. After inoculation with each spore suspension, the sets were placed in a dew chamber for 24 h at 15°C and then in a growth chamber maintained at a temperature between 18 and 22°C with a 16-h photoperiod (daylight supplemented with 400 W sodium lamps). The infection type on the differential lines was scored visually 11 days after inoculation, based on the 0-to-4 scale described by Stakman et al. (1962). Each isolate was assigned a seven-digit code adapted from the scoring system proposed by Gilmour (1973). The 20 lines of the differential set (18 NILs, ‘CS2A/2M’ and ‘Harrier’) were arranged in six sets of three lines, and one set of two *Lr* genes, with each tested isolate assigned an octal pathotype code.

### Postulation of resistance genes present in the major wheat cultivars

We postulated the *Lr* genes present in the 35 major French wheat cultivars (Table 1) by performing multipathotype tests in a growth chamber under the conditions described above for pathotype determination. The set of standard *P. triticina* isolates used was changed over time, according to the availability of new virulence combinations, to improve the reliability of postulations. Over the 2006-2016 period, this set consisted of 24 isolates (Table S1). A spore suspension (3 mg of spores per ml of mineral oil) was sprayed onto the first leaves of 12 seedlings of each cultivar, with 0.7 ml suspension applied per tray of 20 cultivars. We included a set of 23 differential cultivars in each test, to check the identity and purity of each isolate and the infection type. Infection types were scored with the scale described by Stakman et al. (1962).

### Data analysis

The diversity of virulence phenotypes in the population was estimated with the Shannon-Weaver index (H’):

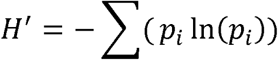

where *p* is the frequency of the *i*^th^ phenotype.

As H’ depends on phenotype frequency, and also on the number of phenotypes (richness), confidence intervals were calculated with the Jackknife procedure of the Species-Richness Prediction and Diversity Estimation packages of R software (Zahl 1977). Evenness was assessed by calculating the *E*^5^ index, also known as the modified Hill’s ratio (i.e. the ratio of the number of abundant phenotypes to the number of rarer phenotypes):

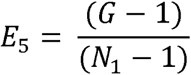

with 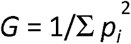 and *N*_1_= *e* ^*H’*^.

The Stoddart and Taylor index *G* represents the number of abundant phenotypes and *N*_1_ represents the number of rarer phenotypes. This index is less dependent on phenotype richness than other evenness indices. E_5_ approaches 0 as a single phenotype begins to predominate (Alatalo 1981).

We first estimated the correlation between the frequency of five of the most abundant pathotypes over the entire 2006-2016 period and their frequency on each of eight of the most widely grown French wheat cultivars during the same period, by calculating Pearson’s correlation coefficient:

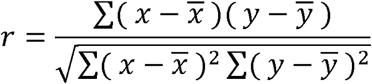

where *x* represents the pathotype frequency in the landscape and *y* its frequency on the eight most widely grown cultivars.

We then performed a principal component analysis (PCA) for each year from 2006 to 2016, for: (i) major cultivars, defined as a cultivar accounting for more than 2% of the pathogen samples in a given year, i.e. 19 of the 35 cultivars presented in Table 2, and (ii) the most abundant pathotypes, defined as pathotypes with a frequency on the 35 major cultivars exceeding 2% in a given year. The cultivar ‘Apache’ was not included in this analysis, because it was considered to display “neutral” behavior, based on the results of the previous overall correlation analysis (see the Results and Discussion sections). The PCA of the association between these two components was visualized with the FactoMinerR package of R software.

## RESULTS

### Diversity of the *P. triticina* population

Two pathotypes dominated the 2006-2016 period in France: 106 314 0 and 166 317 0 (Figure 1). From 2006 to 2015, 106 314 0 was the most frequent pathotype, increasing from 28% in 2006 to 51% in 2009, and then decreasing slightly in frequency to reach a plateau at 30-33% from 2011 to 2014. In 2016, its frequency declined sharply to less than 5%. The other dominant pathotype, 166 317 0, appeared in the landscape in 2007 at a very low frequency (1.4%), then gradually increased to peak at a frequency of 32% in 2012. From 2013 to 2015, the frequency of 166 317 0 decreased to 13%, before increasing to 28% in 2016. Moreover, in 2015, two new pathotypes virulent against *Lr28* appeared (Figure 1): 106 314 2 which had an initial frequency of 1% in 2015, increasing to 14% in 2016; and 167 337 3, which had an initial frequency of 12% in 2015, staying at 10% in 2016. Taken together, these two pathotypes accounted for 24% of the total *P. triticina* population in 2016.

**Figure 1.**
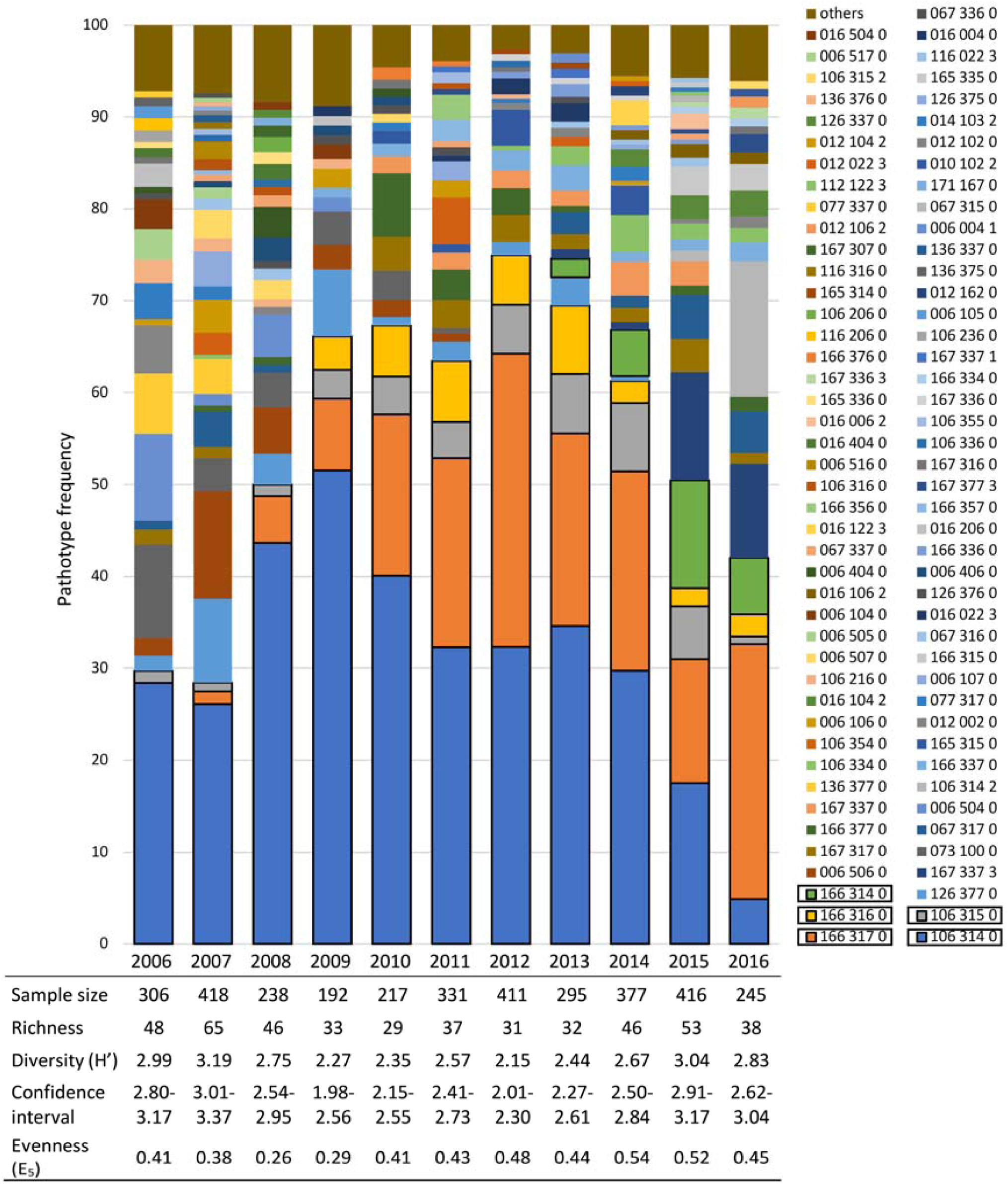
Frequency of *Puccinia triticina* pathotypes during the 2006-2016 period and associated indices of richness, diversity (H’), and evenness (E_5_). The confidence interval for the Shannon-Weaver diversity index (H’) was calculated with the Jackknife procedure. Pathotypes were characterized on a differential set of 20-*Lr* genes. Five of the most prevalent pathotypes during the period, including 106 314 0 and 166 317 0, are boxed. “Others” corresponds to pathotypes found only once in the year.

With the exception of the four abovementioned pathotypes, the frequency of individual pathotypes never exceeded 11% annually. The overall proportion of all other pathotypes decreased from 73% in 2006 and 2007 to 36% in 2012, before increasing again to 50% over the last two years of the time period studied. This overall trend was formalized by a decrease in pathotype diversity from 2009 to 2014, with richness (*n* = 31) and Shannon-Weaver index (H’ = 2.15) values lowest in 2012 (Figure 1), subsequently increasing until 2015 (richness *n* = 53 and H’= 3.04), before declining again in 2016 (richness = 38 and H’= 2,83). The evenness of pathotype distribution, represented by the E_5_ index, remained stable over the 2006-2016 period (ranging from 0.38 to 0.54), but with a transient decrease in 2008 (0.26) and 2009 (0.29).

### Changes in virulence frequencies in the pathogen population and prevalence of the corresponding resistance *Lr* genes in the varietal landscape

Virulence genes could be categorized into three groups according to the pattern of change in their frequencies in the French *P. triticina* population during the 2006-2016 period. The first group (Figure 2A, represented in blue) included virulence genes against *Lr14a, Lr10, Lr13, Lr37, Lr17, Lr15, Lr1*, which frequency in the pathogen population increased rapidly to more than 80%, from 2009 to 2016. In terms of the frequency of the corresponding *Lr* genes in the host population (Figure 2B), *Lr13* and *Lr37* were the most represented, with a frequency above 56%, peaking at 72% for *Lr13* in 2006. The frequency of *Lr14a* was initially 16% in 2006, and increased to 46% in 2016. The frequency of *Lr1* was stable at 6% over the 11 years. *Lr10* was the only resistance gene for which a decrease in frequency was observed, from 40% in 2006 to 15% in 2016.

**Figure 2.**
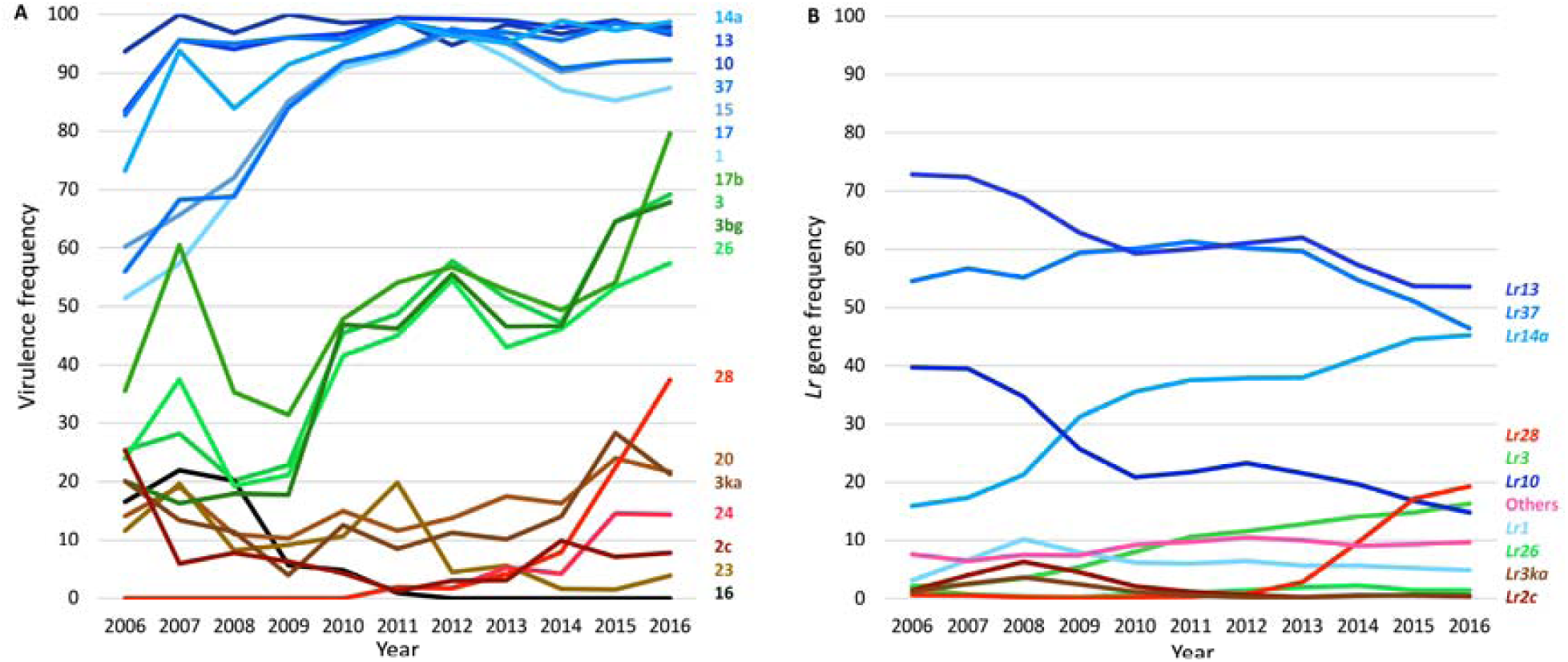
Changes in virulence frequencies in the *Puccinia triticina* population (A) and in the frequencies of the corresponding *Lr* resistance genes in the host population (B) in the French landscape during the 2006-2016 period. Virulence phenotypes were identified on a differential set of 20-*Lr* genes, and *Lr* gene combinations in the cultivars were postulated based on a set of standard isolates. “Others” corresponds to cultivar mixtures, unidentified cultivars and cultivars carrying unidentified genes.

The second group (Figure 2A, represented in green) included virulence genes against *Lr17b, Lr3, Lr3bg*, and *Lr26*, with a first peak in frequency at the start of the study period (e.g. virulence 17b at 60% in 2007), followed by a transient decrease and then a gradual increase in frequency to between 57% (virulence 26) and 80% (virulence 17b) in 2016. Two of these four genes, *Lr17b* and *Lr3bg*, were never postulated in the cultivars, whereas the frequency of *Lr26* remained stable over the entire period, at less than 3%, and that of *Lr3* increased steadily, from 1% in 2006 to 16% in 2016 (Figure 2B), making *Lr3* the fifth most frequent *Lr* gene in 2016.

The third group (Figure 2A, represented in brown) comprised four virulence genes against *Lr20, Lr3ka, Lr2c* and *Lr23*, with frequencies that remained below 30%. Of the four corresponding *Lr* genes, only *Lr3ka* and *Lr2c* were postulated in the cultivars, at a very low frequency: less than 7% for *Lr2c* and 3% for *Lr3ka* (Figure2B).

Virulence against *Lr16*, which had a frequency of about 16% at the start of the study period (Figure 2A), was no longer detected after 2011, and *Lr16* was never postulated in the cultivars.

New virulences emerged in 2011 (Figure 2A, represented in red), increasing in frequency until 2016, to values of 15% for virulence 24 and 37% for virulence 28. The frequency of *Lr24* in the cultivars remained very low, at less than 1% (data not shown). The frequency of *Lr28* in the cultivars increased steadily, from 1% in 2012 to 19% at the end of the study period (Figure 2B, represented in red).

Part of the landscape, consistently about 10% over the 2006-2016 period, was planted with cultivar mixtures, unidentified cultivars, or cultivars carrying unidentified genes (Figure 2B).

### Pattern of association between *P. triticina* pathotypes and wheat cultivars

The frequency of four of the five most frequent pathotypes in the landscape (106 314 0, 106 315 0, 166 314 0 and 166 317 0) was correlated with their frequency on ‘Apache’ over the entire 2006-2016 period (*p* < 0.01; Table 3). ‘Apache’ was the only cultivar for which this was the case over the whole period. This result justified a PCA excluding ‘Apache’, which was considered to display “neutral” behavior (see Discussion section), to make it possible to detect associations.

**Table 3.** Pearson’s correlation coefficient for the relationship between the frequency of the five most prevalent pathotypes in the landscape during the 2006-2016 period, and their frequency on eight major cultivars

| Pathotype | Cultivar |  |  |  |  |  |  |  |
| --- | --- | --- | --- | --- | --- | --- | --- | --- |
|  | Arezzo | Apache | Expert | Aubusson | Premio | Bermude | Altigo | Cellule |
| 106 314 0 | **0.88\*** | **0.86\*** | 0.76 | 0.16 | **1.00\*** | 0.57 | - | - |
| 106 315 0 | 0.57 | **0.90\*** | -0.53 | 0.44 | -0.48 | **0.93\*** | - | - |
| 166 314 0 | 0.87 | **0.92\*** | 0.87 | - | - | - | 0.78 | 1.00 |
| 166 316 0 | 0.12 | 0.67 | 0.70 | 0.73 | 0.46 | -0.23 | 0.96 | 1.00 |
| 166 317 0 | 0.61 | **0.82\*** | 0.89 | 0.85 | 0.98 | 0.66 | 0.69 | 0.85 |

\*Significant coefficients ( $p$ -value < 0.01) are shown in bold typeface.
- Pathotype never found on the cultivar

The frequency in the landscape of pathotype 106 314 0, which is avirulent on cultivars ‘Altigo’ and ‘Cellule’ carrying *Lr3*, was significantly correlated with its frequency on cultivars ‘Arezzo’, ‘Apache’ and ‘Premio’, but not with its frequency on ‘Expert’, ‘Aubusson’ or ‘Bermude’, over the entire 2006-2016 period (Table 3). The PCA refined the association between the major cultivars and the most abundant pathotypes for each year of the 2006-2016 period separately, and, thus, independently of changes in the pathogen population that could potentially alter or hide such a relationship in an overall analysis of the whole dataset. The first dimension of the PCA explained at least 48.9% and, at most, 79.1% of the variability, depending on the year considered (Figure 3). Based on the PCA results, we distinguished a group of cultivars associated with pathotype 106 314 0 from a group of cultivars associated with 166 317 0. Pathotype 106 314 0 was always associated with several cultivars, the precise identity of which depended on the year: ‘Aubusson’, ‘Tremie’, ‘Sankara’ and ‘Charger’ from 2006 to 2009, then ‘Arezzo’ in 2010, 2011, and subsequently ‘Bermude’ and ‘Aubusson’ until 2015. In 2015, new associations with cultivars ‘Pakito’, ‘Solehio’ and ‘Expert’ were highlighted. In 2016, 106 314 0 was weakly associated with only one cultivar, ‘Expert’.

**Figure 3.**
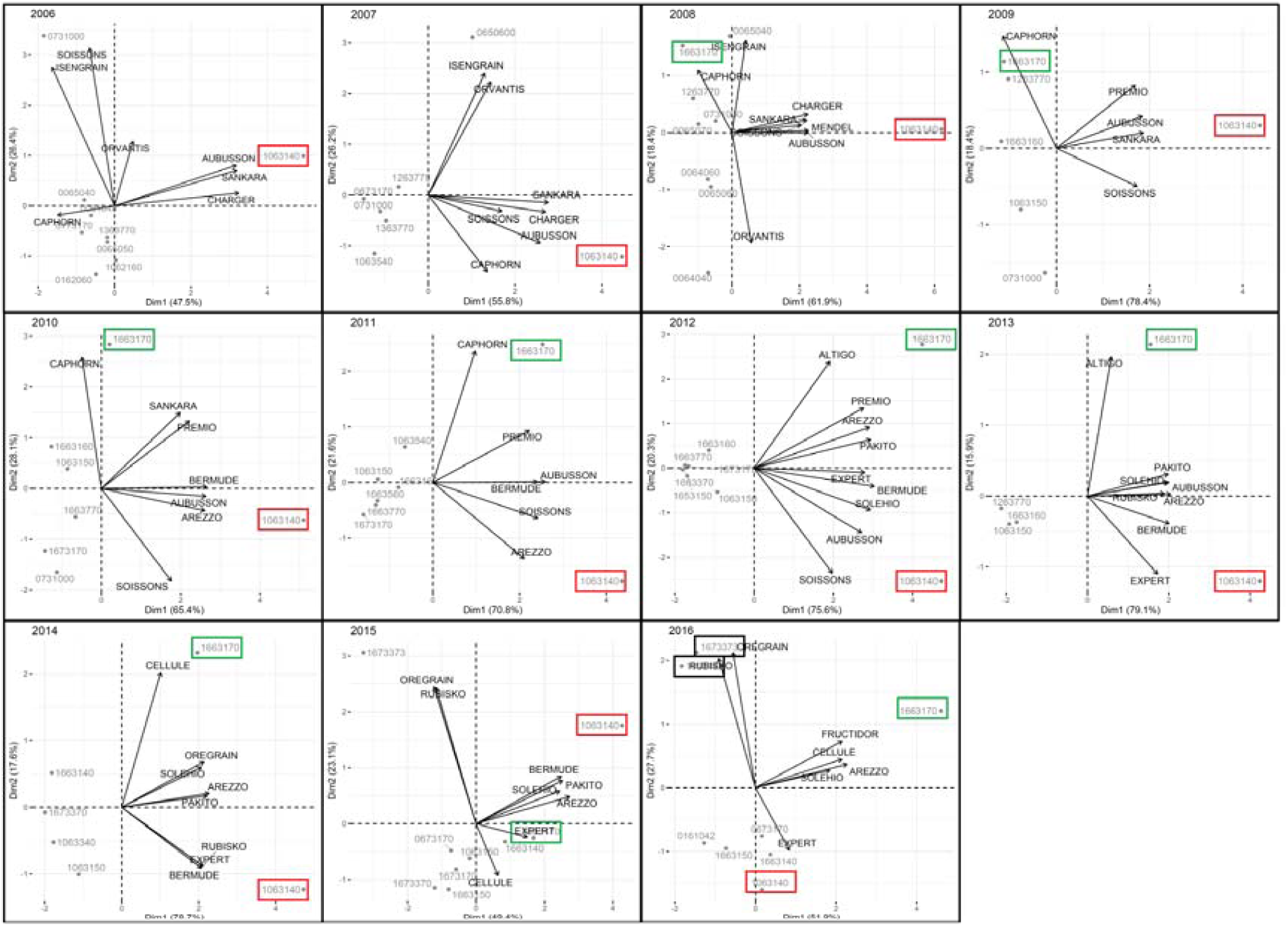
PCA representing 19 of the 35 most widely grown cultivars (Table 1) as variables and *Puccinia triticina* pathotypes as individuals for each year between 2006 and 2016. The two most prevalent pathotypes, 106 314 0 and 166 317 0, are boxed in red and green, respectively. The two pathotypes that emerged in 2015, 167 337 3 and 106 314 2, are boxed in black

The frequency of pathotype 166 317 0 in the landscape was correlated with its frequency on only one cultivar, ‘Apache’, over the entire 2006-2016 period (Table 3). PCA showed that pathotype 166 317 0 was mostly associated with a single cultivar in each year (Figure 3), but that this cultivar differed between years during the 2006-2016 period: ‘Caphorn’ from 2008 to 2011, then ‘Altigo’ in 2012-2013, and finally ‘Cellule’ in 2014-2015. In 2016, unlike other years, this pathotype was associated with several cultivars, ‘Cellule’, ‘Fructidor’, ‘Arezzo’ and ‘Solehio’.

The 167 337 3 and 106 314 2 pathotypes emerged in 2015 and appeared to be strongly associated with new cultivars carrying the *Lr28* resistance gene — ‘Oregrain’, ‘Rubisko’ and ‘Nemo’ — in 2015 and 2016 (Figure 3).

## DISCUSSION

In this study, we analyzed the changes in virulence phenotypes in the French *P. triticina* population over an 11-year period. We found that changes in pathotype distribution depended on the *Lr* genes present in the cultivar landscape, consistent with previous findings. However, we also found that *Lr* genes were not sufficient to account fully for the observed pathotype distribution. This conclusion, which may appear counterintuitive, is robust, given the extensive nature of our survey: samples of *P. triticina* were collected from the most common wheat cultivars in a network of about 50 nurseries distributed throughout the main wheat-growing areas in France. The diversity and pathotype frequency found in such nurseries are known to reflect the situation in the commercial wheat-field landscape (Kolmer 1992).

### The Lr genes present in the varietal landscape are an essential driver of the evolution of P. triticina populations

Changes in the frequency of virulence phenotypes in the *P. triticina* population appeared to depend on the changes in and prevalence of *Lr* genes in the varietal landscape. An analysis of the distribution of virulence phenotype (pathotype) frequencies revealed that one pathotype, 106 314 0, virulent against *Lr1, Lr10, Lr13, Lr14a, Lr15, Lr17* and *Lr37*, predominated in France from 2006 to 2016. Three of these virulences matched the most prevalent *Lr* genes in the French landscape: *Lr13, Lr14a* and *Lr37*. This finding suggests that this virulent pathotype was subject to active selection due to the presence of these three *Lr* genes in the cultivars and, thus, the frequency of these cultivars in the landscape. It has already been shown that *Lr* genes drive the selection of virulence phenotypes. For instance, *Lr39* was widely used in the US, leading to the selection of *P. triticina* populations in which more than 50% of the strains were virulent against this gene (Kolmer 2019). Older datasets available in France suggest that the *Lr* genes present in the varietal landscape drive the selection of virulence phenotypes; from 1999 to 2002, *Lr13* and *Lr14a* were the two most frequent resistance genes present in the cultivars grown, and corresponding virulences were the two most common in the *P. triticina* population (Goyeau et al. 2006).

The group of pathotypes carrying virulence against *Lr3* steadily increased in frequency over the 2006-2016 period. This may reflect the strong pressure imposed by the increasing frequency of the *Lr3* resistance gene in the varietal landscape. As this virulence was carried by pathotype 166 317 0, but not by 106 314 0, the presence of *Lr3* in the landscape probably conferred an advantage on 166 317 0. This finding is consistent with the outputs of the PCA for 2012 data, which highlighted an association of pathotype 166 317 0 with cultivars ‘Altigo’ and ‘Cellule’ carrying *Lr3. Lr3* was deployed for the first time in France in two cultivars registered in 1988, ‘Génial’ and ‘Louvre’, and subsequently in six other cultivars registered in 1998 (Goyeau & Lannou 2011). This deployment of cultivars carrying *Lr3* resistance was followed by a gradual increase in the frequency of virulent pathotypes, including 166 317 0. New virulences against *Lr24* and *Lr28* appeared in 2011. *Lr24* had been present in the varietal landscape since 2006, but at a very low frequency, remaining below 1% until 2012. The increase in the frequency of virulence against *Lr28* immediately followed the registration of cultivars carrying *Lr28*, but began before the widespread deployment of these cultivars (<1% in 2011 and 2012; >9 % in 2014), suggesting that this resistance gene acted as a strong selective driver of the evolution of the *P. triticina* population. *Lr28* resistance was overcome within two years (2014-2015), with a large decrease of its efficacy in France. A similar pattern was observed in the Czech Republic, where *Lr28* conferred high levels of resistance (rated 7.0) to cultivars widely deployed in 2013, but displayed a rapid loss of efficacy, with resistance levels falling to 2.7 for these cultivars in 2015 (Hanzalová et al. 2021). *Lr* genes are generally rapidly overcome in wheat-growing areas worldwide. For example, *Lr3ka, Lr11* and *Lr24* were overcome within two years of their introduction in North American cultivars (Kolmer 1994, 1996).

Some virulences, such as those directed against *Lr15* and *Lr16*, were detected in the *P. triticina* populations sampled despite the absence of the corresponding *Lr* genes from the cultivars used during the 2006-2016 period. These examples illustrate the general conclusion recently drawn by Kolmer (2019) that resistance genes select strains with specific virulence, but that this virulence may already exist in the pathogen. The indirect selection, or at least an absence of counterselection, of these virulence phenotypes considered “unnecessary” for the pathogen, may come under a “hitchhiking effect” (de Vallavieille-Pope et al. 2011; Persoons et al. 2017).

This survey clearly showed that the frequency of pathotypes is strongly influenced by *Lr* gene prevalence in the French varietal landscape. However, our extensive dataset analysis could not fully account for the observed patterns within the pathogen population, suggesting that there is selection pressure on traits other than virulence.

### *Lr* genes cannot fully account for the frequency distribution of *P. triticina* pathotypes

The frequencies of four of the five most prevalent pathotypes were correlated with the frequency of ‘Apache’, which was the only cultivar involved in a correlation with more than one pathotype. The overall prevalence of these four pathotypes in the French varietal landscape was, thus, quite similar to their frequency on ‘Apache’. This is consistent with the previous finding that no pathotype was preferentially associated with ‘Apache’ in French wheat-growing areas generally (Papaïx et al. 2011). Unlike the other cultivars considered, ‘Apache’ did not exert the significant selective pressure which could be expected from the *Lr* genes it carries. This cultivar was, thus, considered “neutral” and was not included in the PCA.

Two pathotypes predominated in France during the 2006-2016 period. First, 106 314 0 alone, then 106 314 0 in co-dominance with 166 317 0, and finally, 166 317 0 alone. Overall population diversity decreased when these two pathotypes were co-dominant in the varietal landscape (together accounting for more than half the sampled isolates). These dynamics for the 2006 to 2016 period differ from those in France between 1999 and 2002 described in a previous study (Goyeau et al. 2006), suggesting that *P. triticina* diversity is greater when there are two co-dominant pathotypes rather than a single dominant pathotype.

PCA revealed that pathotype 106 314 0 was associated with different cultivars (10 in total) each year from 2006 to 2014. This pattern reflects “generalist” behavior, or, in other words, an adaptation of this pathotype to several host genetic backgrounds. Other pathotypes carrying virulence genes theoretically enabling them to attack the same cultivars were identified through the annual survey, but were never present at a meaningful frequency. For instance, while pathotype 166 317 0 predominated at the end of the 2006-2016 period, other pathotypes virulent against *Lr*3 were found at a much lower frequency on cultivars carrying *Lr3*. The higher prevalence of 106 314 0 and 166 317 0 cannot, therefore, be explained exclusively by their virulence phenotype. Biases due to the focusing of sampling on specific cultivars (the most widely grown) may partly account for such effects, but are unlikely to provide a full explanation. Thus, the selective pressure exerted by *Lr* genes is not the only evolutionary force driving the dynamics of the adaptation of *P. triticina* populations to their host populations at large spatial scales.

### The potential impact of aggressiveness as a driver of changes in pathotype prevalence

A previous study, over the 1999-2008 period in France, demonstrated that some *P. triticina* pathotypes were preferentially associated with certain cultivars whilst being “compatible” with most of the cultivars deployed in the landscape (Papaïx et al. 2011). In this previous study, the resistance level of a cultivar was linked to the frequency of the most aggressive pathotype of all the compatible pathotypes present in the *P. triticina* population. During the period considered by this previous study, the rust population was dominated by one pathotype, 073 100 0, associated with the wheat cultivar ‘Soissons’, which was the most widely grown cultivar in France until 1999 (reaching a maximum of 40% of the total area under wheat in 1993). Pathotype 073 100 0 was more aggressive on ‘Soissons’ than other common virulent pathotypes able to infect this cultivar. This difference in aggressiveness, characterized by a larger uredinium size and higher level of spore production per square millimeter of sporulating tissue, explained the preferential association between Soissons and this pathotype (Pariaud et al. 2009b). Accordingly, as ‘Soissons’ was the most widely grown cultivar, the high frequency of the pathotype 073 100 0 on this cultivar was considered likely to account for its overall prevalence at the landscape scale.

Moreover, Papaïx et al. (2011) highlighted an association of pathotype 106 314 0 with the wheat cultivar ‘Caphorn’, even though this pathotype is considered avirulent on this cultivar at the adult plant stage. During a severe epidemic, pustules of a non-compatible pathotype can, indeed, be found on a cultivar. We did not consider interactions of this type, between incompatible pathotypes and cultivars, in this study.

At the start of this century, the 10 most widely grown cultivars in France accounted for about 70% of the total area under wheat. This percentage dropped between 43 and 56% during the 2006-2016 period (Table 1). The varietal landscape has, thus, tended to become more diversified, potentially accounting for the predominance of a more “generalist” pathotype. The generalist behavior of such a pathotype could be related to its aggressiveness, the quantitative component of pathogenicity (Pariaud et al. 2009a; Lannou 2012). Aggressiveness is considered less dependent on host genetic background. Thus, a *P. triticina* pathotype with a higher aggressiveness on a range of cultivars would be fitter in a diversified or heterogeneous biotic environment (Wilson & Yoshimura 1994; Kröner et al. 2017). The association between pathotype 106 314 0 and the most widely grown cultivars was not related to the compatibility of its virulence profile with the combination of *Lr* genes in these cultivars. It therefore seems likely that aggressiveness traits (i.e. latent period, sporulating capacity and infection efficiency) played a significant role. The selection of groups of strains with higher aggressiveness, as a mechanism driving the evolution of pathogen populations, has been demonstrated in other pathosystems, through empirical evidence of “local adaptation” (Milus et al. 2009; Delmas et al. 2016; Kröner et al. 2017). For instance, in the *Phytophthora infestans* population, a detached-leaflets assay showed that the most aggressive genotypes on a potato cultivar (genotype with the shortest latency period or highest infection efficiency) tended to be those selected by the same cultivar in the field. Moreover, differences in aggressiveness between isolates were amplified on cultivars with the highest levels of partial resistance (Young et al. 2018). In *Plasmopora viticola*, strains collected from resistant grapevine cultivars were more aggressive than isolates collected from susceptible hosts, demonstrating the occurrence of selection for greater aggressiveness (Delmas et al. 2016).

The hypothesis of “dual selection” based on both qualitative and quantitative interactions between *P. triticina* and bread wheat populations, is consistent with the dynamics described here. Unlike pathotype 106 314 0, pathotype 166 317 0 was associated with only one cultivar each year until 2015, first with ‘Caphorn’, then with ‘Altigo’. Pathotype 166 317 0 is virulent against *Lr3, Lr10, Lr13, Lr37*, and would, therefore, theoretically be able to overcome the *Lr* genes carried by ‘Caphorn’ and ‘Altigo’. However, these two cultivars have high levels of resistance to leaf rust (and, indeed, the leaves sampled from these two cultivars carried only a few pustules). We therefore hypothesize that the isolates found on these two cultivars had undergone selection for greater aggressiveness (relative to other strains, when expressed on susceptible cultivars) and that the selective dynamics had turned to their advantage not only at the field scale, but also at the landscape scale. Further experimental studies based on tests of local adaptation (isolate x cultivar cross-inoculations) by comparing aggressiveness traits would be useful to validate this hypothesis and to quantify its potential effects on population evolutionary dynamics. The neutral status of ‘Apache’ renders this cultivar particularly interesting for comparisons of pathotype aggressiveness without a selection effect of the tested cultivar.

## Acknowledgments

We thank Nicolas Lecutier (INRAE BIOGER, Thiverval-Grignon, FR) for technical assistance in the growing of plant material and Dr. Florence Dubs (INRAE UMR GQE-Le Moulon, Gif-sur-Yvette, FR) and Dr. Anne-Lise Boixel (INRAE BIOGER, Thiverval-Grignon, FR) for discussions on statistical analyses. We thank the French Wheat Breeders groups ‘Recherches Génétiques Céréales’ and ‘CETAC’, and ‘ARVALIS-Institut du Végétal’ for their help collecting strains of *P. triticina* in their nurseries and field trials. We thank Dr. Julie Sappa for her help correcting our English.

## Funding

This research was supported by a PhD fellowship from the INRAE department ‘Santé des Plantes et Environnement’ (SPE) and from the French Ministry of Education and Research (MESRI) awarded to Cécilia Fontyn for the 2018-2022 period. It was also supported by several French FSOV (‘Fonds de Soutien à l’Obtention Végétale’) grants (FSOV 2004, 2008, 2012) and by the European Commission, Research and Innovation under the Horizon 2020 program (RUSTWATCH 2018-2022, Grant Agreement no. 773311-2).

## Conflict of Interest Statement

The authors declare that the research was conducted in the absence of any commercial or financial relationships that could be construed as a potential conflict of interest.

---

**Table S1.** Standard isolates of *Puccinia triticina* used in multipathotype tests for the postulation of resistance genes

| Isolate | Pathotype<sup>a</sup> | Virulence expressed at the seedling stage<sup>b</sup> |
| --- | --- | --- |
| B9201-2C3 | 014 103 0 | *Lr2c, Lr13, Lr14a, Lr17b, Lr26* |
| B9407-1CA3 | 050 124 0 | *Lr2c, Lr3bg, Lr14a, Lr20, Lr37* |
| B9833-B | 077 317 0 | *Lr2c, Lr3, Lr3bg, Lr3ka, Lr10, Lr13, Lr14a, Lr15, Lr17, Lr17b, Lr26, Lr37* |
| FSA9 | 150 104 0 | *Lr1, Lr2c, Lr3bg, Lr14a, Lr37* |
| B9506-2B | 014 103 0 | *Lr2c, Lr13, Lr14a, Lr17b, Lr26* |
| B950506A | 106 232 0 | *Lr1, Lr10, Lr13, Lr15, Lr17, Lr17b, Lr20* |
| B950019-A | 012 026 0 | *Lr2c, Lr10, Lr17b, Lr20, Lr37* |
| B950365-D | 073 100 0 | *Lr2c, Lr3, Lr3bg, Lr3ka, Lr10, Lr14a* |
| B01M196-D | 016 206 0 | *Lr2c, Lr10, Lr13, Lr15, Lr17b, Lr37* |
| B03M041 | 116 226 0 | *Lr1, Lr2c, Lr10, Lr13, Lr15, Lr20, Lr17b, Lr37* |
| B03M007 | 077 310 0 | *Lr2c, Lr3, Lr3bg, Lr3ka, Lr10, Lr13, Lr14a, Lr15, Lr17* |
| B00M057-A | 166 336 0 | *Lr1, Lr3, Lr3bg, Lr10, Lr13, Lr14a, Lr15, Lr17, Lr17b, Lr20, Lr37, Lr27 + Lr31* |
| B9384-1C1 | 075 337 0 | *Lr2c, Lr3, Lr3bg, Lr3ka, Lr13, Lr14a, Lr15, Lr17, Lr17b, Lr20, Lr26, Lr37* |
| FS01-D17 | 016 100 0 | *Lr2c, Lr10, Lr13, Lr14a* |
| B77SABA | 775 237 0 | *Lr1, Lr2a, Lr2b, Lr2c, Lr3, Lr3bg, Lr3ka, Lr13, Lr15, Lr17, Lr17b, Lr20, Lr26, Lr37* |
| BT06M140 | 126 377 0 | *Lr1, Lr3, Lr10, Lr13, Lr14a, Lr15, Lr17, Lr17b, Lr20, Lr23, Lr26, Lr37, Lr27 + Lr31* |
| BT08M111 | 775 715 0 | *Lr1, Lr2a, Lr2b, Lr2c, Lr3, Lr3bg, Lr3ka, Lr13, Lr14a, Lr15, Lr16, Lr17, Lr26, Lr37* |
| BT09M168 | 167 337 1 | *Lr1, Lr3, Lr3bg, Lr3ka, Lr10, Lr13, Lr14a, Lr15, Lr17, Lr17b, Lr20, Lr24, Lr26, Lr37, Lr27 + Lr31* |
| BT11M182-A | 016 022 2 | *Lr2c, Lr10, Lr13, Lr17b, Lr20, Lr28* |
| BT12M097 | 012 022 3 | *Lr2c, Lr10, Lr17b, Lr20, Lr24, Lr28* |
| BT12M413-3 | 002 004 2 | *Lr10, Lr28, Lr37* |
| BT14M228 | 016 122 3 | *Lr2c, Lr10, Lr13, Lr14a, Lr17b, Lr20, Lr24, Lr28* |
| BT16M051 | 167 337 3 | *Lr1, Lr3, Lr3bg, Lr3ka, Lr10, Lr13, Lr14a, Lr15, Lr17, Lr17b, Lr20, Lr24, Lr26, Lr28, Lr37, Lr27 + Lr31* |
| BT16M225 | 012 106 2 | *Lr2c, Lr10, Lr14a, Lr17b, Lr28, Lr37* |

<sup>a</sup> Seven-digit triplet code (Gilmour, 1973) based on a 20-*Lr* gene differential set arranged as follows: [*Lr1, Lr2a, Lr2b*], [*Lr2c, Lr3, Lr3bg*], [*Lr3ka, Lr10, Lr13*], [*Lr14a, Lr15, Lr16*], [*Lr17, Lr20, Lr23*], [*Lr26, Lr17b* (the Australian cv. 'Harrier'), *Lr37*] and [*Lr24, , Lr28* (line CS2A/2M)].
<sup>b</sup> Tested for pathogenicity on the 18 Thatcher NILs, the Australian cv. Harrier (*Lr17b*), CS2A/2M (*Lr28*), the lines with the resistance genes *Lr9, Lr19*, and the Australian cv. Gatcher (*Lr10, Lr27+Lr31*).

**Table S2.**
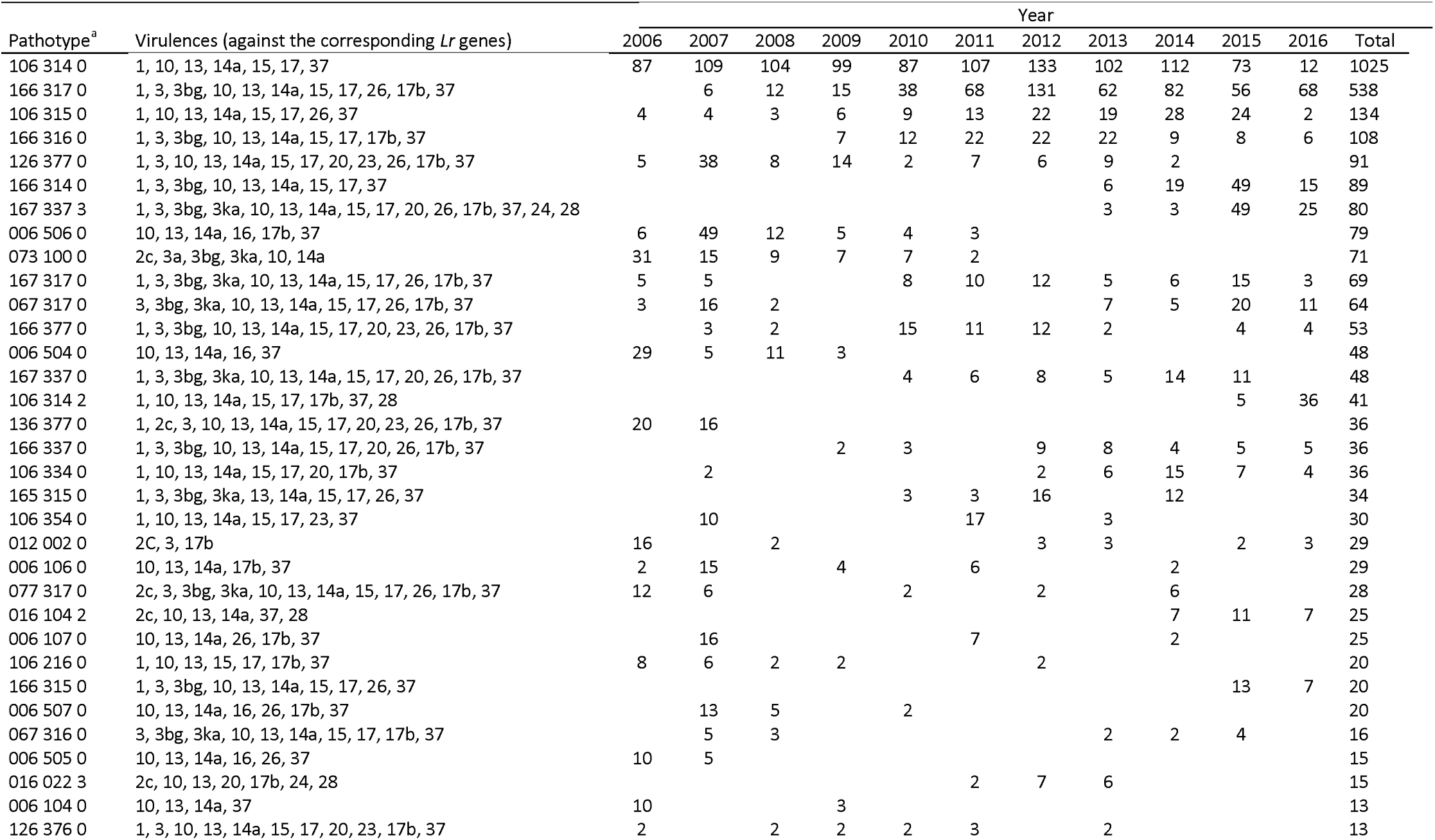
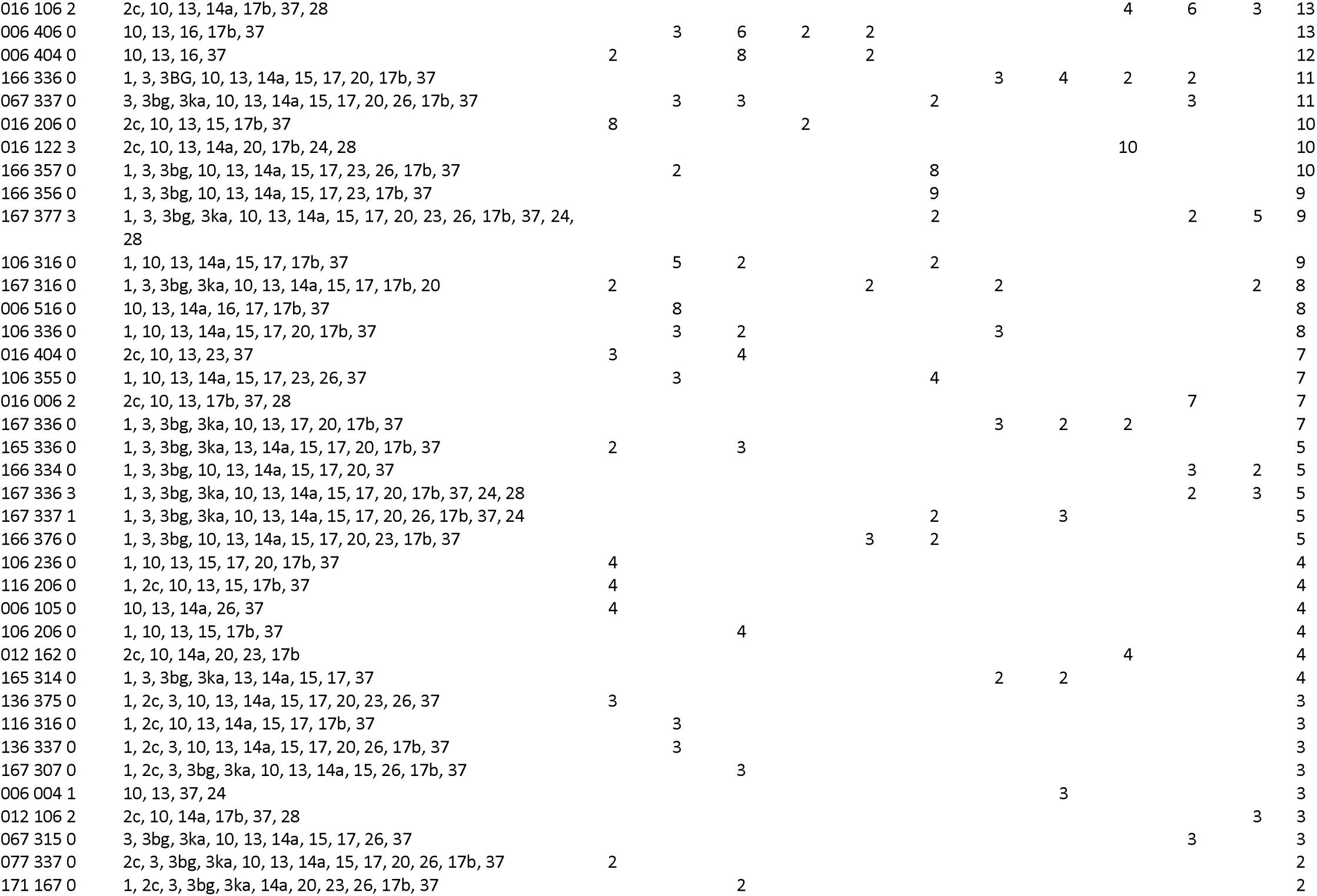
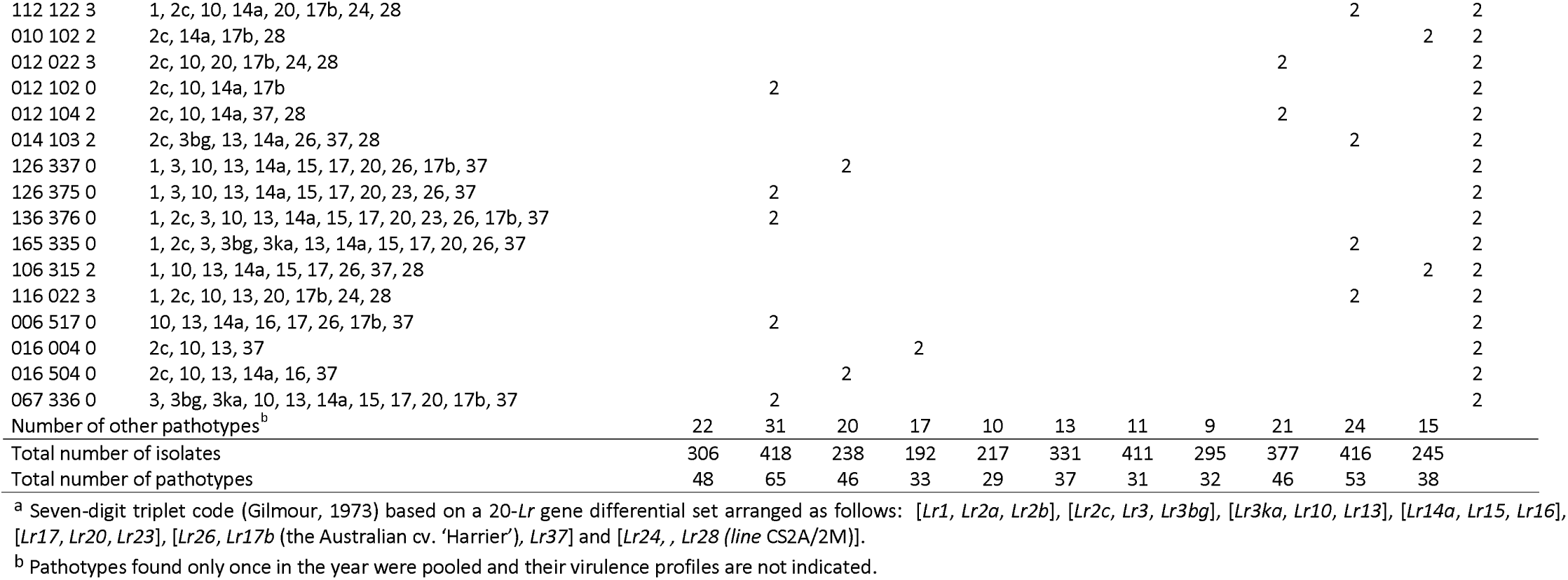
Number of *Puccinia triticina* isolates collected from wheat during the period 2006-2016, by pathotype, as determined with a differential set of 20 *Lr* genes

## Notes

### Competing Interest Statement

The authors have declared no competing interest.

